# A transcriptomic-driven segmentation and cell simulation framework for high-resolution spatial transcriptomics and cell-cell communication

**DOI:** 10.64898/2026.04.24.720489

**Authors:** Visanu Wanchai, Nancy C Bustamante-Gomez, Alongkorn Kurilung, Karen E. Beenken, Sergio Cortes, Mark S. Smeltzer, Yuet-Kin Leung, Jinhu Xiong, Maria Almeida, Charles A O’Brien, Intawat Nookaew

## Abstract

High-resolution spatial transcriptomics platforms, such as Visium HD and Atera, enable transcriptome-wide profiling at near-single-cell resolution. However, accurate cell segmentation to define spatial boundaries relies heavily on histological images. Previous approaches struggle in tissues with high cell density, inflammation, or mineralization, leading to transcriptomic bleed-through and inaccurate clustering. To address this, we developed TENGU (<u>T</u>ranscript-signal <u>E</u>nrichment and <u>G</u>rouping <u>U</u>nit), a comprehensive end-to-end bioinformatic software package. Unlike existing tools, TENGU employs a transcript-first segmentation approach, prioritizing transcript-signal density as the primary modality and utilizing histological images only as a secondary supplement in unresolved regions. These initial boundaries are further optimized through a transcriptomic-driven cell simulation algorithm that iteratively refines boundaries based on localized gene expression probabilities, effectively minimizing spatial scattering and preserving biologically distinct molecular signatures. The pipeline seamlessly integrates tissue segmentation, high-resolution cell-type annotation, and spatially aware cell-cell communication (CCC) analysis. We rigorously benchmarked TENGU against the 10x Genomics and Bin2cell pipelines for segmentation, and against CellNEST for spatial CCC inference, across diverse and technically challenging microenvironments. TENGU demonstrated superior transcriptomic distinctness in the murine brain and successfully captured matrix-embedded osteocytes in decalcified bone. In murine osteomyelitis, TENGU outperformed CellNEST in sensitivity, localizing both reactive bone remodeling (Tgfb) and dense, abscess-encapsulating inflammatory (Il1a/Il1) networks. TENGU also resolved species-specific pro-tumorigenic hubs (MDK-SDC4) in a human colorectal cancer xenograft and generalized effectively to the Atera platform, delineating CXCL12-CXCR4 and LGALS3-TGFBR2 signaling within highly compacted human breast cancer tissue. By mitigating image-dependent constraints, TENGU provides a highly adaptable, robust computational framework to decode the functional micro-anatomy of healthy and pathological tissues.

**Key points:** TENGU delivers three major advancements to the field of spatial biology:

- Transcript-First Segmentation: Rather than relying solely on histology, TENGU prioritizes molecular density to define primary segmentation boundaries, successfully capturing cells that are often missed or obscured in standard imaging.
- Transcriptomic-Driven Cell Simulation: To denoise the data and resolve transcript diffusion, TENGU utilizes a probabilistic cell simulation framework. By iteratively refining cell shapes based on localized gene expression probabilities, this algorithm dynamically groups highly correlated transcripts and restores biologically distinct molecular signatures.
- End-to-End Analytical Pipeline: High-fidelity segmentation is seamlessly integrated into a complete downstream workflow. TENGU provides built-in modules spanning from spatial object embedding and automated, high-resolution cell-type annotation, to spatially aware cell-cell communication (CCC) analysis, delivering actionable functional insights in a single unified package.

## Introduction

The advent of high-resolution spatial transcriptomics, such as the 10x Genomics Visium HD platform, has redefined our capacity to investigate tissue architecture by mapping whole-transcriptome expression at a near single-cell resolution based on a contiguous grid of 2 µm squares [1, 2]. Although this and other platforms have enabled the capture of massive amounts of transcript information, correct assignment of a given transcript to a specific cell, which requires definition of cell boundaries, in the tissue sample remains challenging. Correct assignment of transcripts to their cell of origin is essential for meaningful cell type annotation and analysis of cell to cell communication.

Current computational pipelines (e.g., Space Ranger, Bin2cell, ENACT) attempt to define cell boundaries by relying heavily on morphological assumptions derived from corresponding images of Hematoxylin and Eosin (H&E) stained tissue sections [3–5]. Unfortunately, these image-based approaches often fail to account for biological variation of cell shapes, inconsistent sample quality, and lateral diffusion of transcripts during tissue permeabilization. Consequently, they suffer from the misassignment of transcripts to adjacent cells and on the incorrect identification of cell boundaries, which confounds downstream analysis [6–8].

In this study, we introduce TENGU (Transcript-signal Enrichment and Grouping Unit), a comprehensive bioinformatic software package specifically designed to overcome these limitations for Visium HD data. TENGU advances spatial transcriptomic analysis by delivering three core capabilities. First TENGU uses transcript-signal based segmentation. Rather than relying solely on histology, TENGU prioritizes molecular density to define primary segmentation boundaries, capturing cells that may be missed or obscured in standard imaging. Second, TENGU uses transcriptomic-driven cell simulation. To denoise the data and resolve transcript diffusion, TENGU utilizes the probabilistic cell simulation framework, Proseg [9], which requires prior segmentation as the input. By iteratively refining the segmented shapes based on the likelihood of their localized gene expression patterns, this step dynamically groups highly correlated transcripts and restores biologically distinct molecular signatures. Third, TENGU provides an end-to-end analytical pipeline. High-fidelity segmentation is seamlessly integrated into a comprehensive downstream workflow to advance biological interpretation. TENGU features built-in modules for spatial object embedding, interactive visualization exporting, automated high-resolution cell-type annotation, and spatially aware cell-cell communication (CCC) analysis, delivering actionable functional insights within a single, unified package.

We rigorously benchmarked TENGU against the standard 10x Genomics pipeline and Bin2cell. Our results demonstrate TENGU’s superior transcriptomic distinctness and spatial resolution across highly diverse and technically challenging microenvironments, including murine brain, decalcified murine bone, and a highly compacted, dual-species human colorectal cancer xenograft.

## Materials and Methods

### Murine osteomyelitis model

Wild type C57BL/6J mice were obtained from the Jackson Laboratory. Mice were housed at 2-5 animals per cage using a blend of 1/4″ corncob bedding and white enrichment paper both produced by Andersons Incorporated. Mice were provided ad libitum water and an irradiated Purina diet of 5V5R. The temperature range in the room was 68-79 F with a set point of 71 +/-2. Room humidity ranged from 30% to 70%. The room was on a 12:12 hour light cycle and the illumination was 364 lux measured 1 meter from the floor. Mice were anesthetized and the right femur was surgically exposed as previously described [10, 11]. A unicortical defect was generated using a 21-guage needle and the trough was inoculated with 10^6^ colony forming units (CFU) of *S. aureus* USA300 strain LAC suspended in 2 μL PBS. Muscle fasciae and skin were closed and infection was allowed to proceed for 10 days, at which point mice were euthanized and the surgical limb was harvested. All animal procedures were reviewed and approved by Institutional Animal Care and Use Committee of the University of Arkansas for Medical Sciences.

### RNA in situ hybridization

Femurs were fixed in 10% Millonig’s formalin for 48 h at 4 °C and decalcified in 14% EDTA (pH 7.4) for 7 days at 4 °C. Samples were dehydrated in 100% ethanol, embedded in paraffin, and sectioned at 8 µm. RNA *in situ* hybridization was performed using the RNAscope® 2.5 HD Assay-Red according to the manufacturer’s instructions. Briefly, sections were deparaffinized and incubated with 3% hydrogen peroxide for 10 min at room temperature to quench endogenous peroxidase activity. Sections were then treated with RNAscope® Custom Pretreatment Reagent (Advanced Cell Diagnostics, USA) for 30 min to enhance tissue permeability. Hybridization was performed with the mouse *Tnfsf11*(red) and *Acp5*(blue) probe for 2 h at 40 °C in a HybEZ II oven (Advanced Cell Diagnostics, USA). Signal amplification was carried out sequentially using AMP1– AMP6 reagents with wash buffer steps between incubations. Fast Red chromogen was prepared by mixing Fast Red-A and Fast Red-B at a 60:1 ratio and applied for 10 min at room temperature. Sections were counterstained with 25% hematoxylin, blued in 0.02% ammonia water, dried at 60 °C for 5 min, and mounted with EcoMount (Biocare Medical, USA) prior to imaging on a Keyence BZ-X810 microscope (Keyence, USA).

### Visium HD Spatial transcriptomic analysis

Bone tissue sections used for Visium HD were prepared using the same FFPE processing and sectioning procedure described above for RNAscope®. Tissue sections were deparaffinized and decrosslinked according to the Visium HD FFPE Tissue Preparation Handbook (CG000684, Rev D, 10× Genomics, USA). Probe hybridization, ligation, and transfer were performed using the Visium HD Spatial Gene Expression workflow (CG000685, Rev A, 10× Genomics, USA). Briefly, mouse whole-transcriptome probe set v2 was hybridized to tissue sections for 18 h, followed by probe ligation. Ligated probe products were then released and transferred to a Visium HD slide (6.5 × 6.5 mm capture area) using the CytAssist instrument (10× Genomics, USA). Probe extension and amplification were subsequently performed to generate sequencing libraries. Final libraries were indexed using Dual Index TS Set A (10x Genomics) and sequenced on an Illumina NovaSeq X Plus using paired-end, dual-index reads (Read 1: 151 cycles; i7: 10 cycles; i5: 10 cycles; Read 2: 151 cycles).

Preprocessing of Visium HD data (.fastq files) was performed with Space Ranger v4.0.1. While the pipeline provides outputs at the native 2 µm resolution as well as at 8 µm and 16 µm binning, all downstream analyses exclusively utilized the 2 µm-binned data.

### Transcriptomic-driven segmentation and cell simulation

To identify segmented boundaries, the TENGU framework utilizes a two-stage computational pipeline: primary segmentation and subsequent probabilistic refinement. The framework integrates and extends Bin2Cell v0.3.4 [4] and Proseg v3.1.0 [9] to execute robust segmentation inference in Visium HD data. TENGU adapts Bin2Cell into a transcript-signal-first approach, where spatially resolved transcript counts (2 µm resolution) were converted into a continuous signal representation. StarDist v0.9.2 [12] equipped with the 2D versatile fluorescence model, was applied to this signal to infer cell boundaries independent of histology. Optimization was performed on three critical parameters to maximize the median gene count per segmented unit: (i) the Gaussian filtering kernel (sigma) to normalize input signals; (ii) the probability threshold (pt) for cell candidate detection; and (iii) the non-maximum suppression (nms) threshold to regulate adjacent unit overlap. In complex regions with high cellular density or low transcript contrast, this primary segmentation was augmented via a secondary image-based step using the StarDist 2D 10X H&E model on corresponding histological image. Reconciling these transcript-signal- and image-based modalities produced a unified base segmentation map. Finally, Proseg [9] was deployed to perform probabilistic cell simulation on the base segmentation. By modeling spatial and transcriptomic uncertainty, Proseg iteratively adjusts cell geometry and reassigns transcripts, yielding a final set of refined segmented units suitable for advanced downstream analysis (Supplementary Fig S1 for details of the workflow).

### Cell type annotation

The TENGU pipeline includes a dedicated, flexible module for automated cell-type annotation. While the architecture readily supports alternative prediction methods, for this study we utilized CellTypist v1.7.1 [13], a supervised machine-learning classifier. Prior to classification, Visium HD gene expression matrices were log-normalized and formatted to meet CellTypist requirements. Predictions were generated using two categories of reference models. First, we utilized publicly available, pre-trained CellTypist models derived from whole adult mouse brain [14] and human colorectal cancer [15] datasets. Second, we deployed a custom CellTypist model trained on our previously published murine bone dataset [16],which was specifically designed to primarily capture transcriptional heterogeneity within mesenchymal lineages and with additional transcriptional heterogeneity of hematopoietic cells. For all analyses, CellTypist was executed in standard prediction mode, yielding discrete cell-type assignments and their associated confidence scores for each segmented unit.

To ensure broad interoperability, all final segmented objects and cell type annotation can be exported to the open-standard .zarr format for utilization by third-party analytical tools. We use python package SpatialData [17] version 0.6 and R suite software verion 4.5 for data analysis and visualization. Furthermore, the data can be exported in the .loupe format, allowing for interactive spatial visualization of the results via Loupe Browser v9.0.0.

### Ligand-receptor interaction analysis

To quantitatively evaluate spatially constrained cell-cell communication, we derived a distance-weighted ligand-receptor interaction score for all potential sender-receiver cell pairs. Following the application of a baseline expression threshold (positive counts), ligand-expressing (sender) and receptor-expressing (receiver) cells were identified. The spatial proximity between any given ligand-positive cell *i* and receptor-positive cell *j* was determined by calculating the Euclidean distance between their segmentation polygon centroids. The final cell-cell interaction score (CCS) incorporates ligand expression, receptor expression, and an exponential spatial-decay penalty, defined mathematically as:

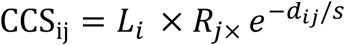

Where, *L_i_* represents the ligand expression level in the sender cell, *R_j_* represents the receptor expression level in the receiver cell, *d_ij_* is the Euclidean distance between the centroids, and *S* is the global median cell-to-cell distance. This exponential decay term ensures that distal interactions are heavily penalized, accurately modeling the reduced probability of functional ligand diffusion across the tissue microenvironment. The results derived from TENGU were compare with the results from CellNEST [18] (default parameters) base on the same ligand-recepter dataset.

### Quantification of Transcriptomic Distinctness

To quantitatively evaluate the transcriptomic distinctness of the segmented cell populations and assess the robustness of our clustering, we calculated the Silhouette score across a range of dimensionality thresholds. To avoid the non-linear distance distortions introduced by 2D projections such as UMAP, this metric was computed directly within the high-dimensional principal component analysis (PCA) space. To ensure computational efficiency and eliminate sampling bias across different evaluations, a consistent random subsample of 10,000 cells was selected using a fixed random seed. The Silhouette score was subsequently calculated on this exact cellular subset across a varying number of top principal components (ranging from 2 to 50) using the scikit-learn library in Python. The optimal dimensionality for evaluating transcriptomic mixing was determined by identifying the peak Silhouette score, which represents the mathematical optimum between capturing true biological variance and minimizing technical noise.

### Quantification of Spatial Scattering

To quantitatively assess the degree of spatial scattering and transcriptomic mixing among the segmented units, we calculated a Neighborhood Purity score for each segmentation method. For every individual cell, we identified its different k (5-1,000) closest physical neighbors in the 2D tissue coordinate space using a K nearest neighbors algorithm implemented via the Python scikit-learn library. The local purity for each cell was defined as the fraction of its k neighbors that shared the exact same cell type label. These local values were then averaged across all cells in the dataset to generate a global Neighborhood Purity score. A global score closer to 1 indicates that cells are primarily surrounded by their own type (forming biologically cohesive, contiguous domains), whereas lower scores indicate high spatial dispersion and transcriptomic mixing.

### Atera spatial transcriptomic analysis by TENGU

To demonstrate TENGU’s versatility with high-resolution (0.2 µm) image-based transcriptomics, we analyzed a genome-wide dataset from the Atera platform (10X genomics). Because Atera captures transcript signals in three dimensions (x-y-z), the raw data was first dimensionally reduced and aggregated into a two-dimensional format (x-y) to match the Visium HD data structure. The preprocessed two-dimensional spatial data was then input into the TENGU pipeline.

## Results

### Comprehensive bioinformatic software package for Visium HD data analysis

The computational workflow of TENGU, summarized in Figure 1A, consists of three major procedures. 1) **Segmentation**: TENGU adapts the Bin2Cell [4] pipeline, utilizing both transcript-and image-based modalities. For the primary, transcript-based approach, TENGU applies the StarDist 2D versatile fluorescence model [12]. We optimized three key parameters to maximize the median number of genes per segmented unit (a common metric to assess spatial transcriptomics data): (i) input signal normalization (by varying Gaussian filtering (sigma) for image-based signal-to- transcript noise reduction), (ii) the probability threshold ((pt) to modulate cell detection from the signal), and (iii) the non-maximum suppression threshold ((nms) to control the degree of overlap among segmented units). The parameter optimization results for representative mouse samples, utilizing either probe-based chemistry (bone and brain) or 3’ poly(A) capture-based chemistry (brain and xenograft), are illustrated in Figure 1B. The same parameter set of sigma = 50, pt = 0.01 and nms = 0.01, yielded the highest number of genes per segmented unit. In dense cellular areas where unresolved signal boundaries hinder this primary approach, segmentation is supplemented by a secondary image-based step utilizing the StarDist 2D 10X H&E model [3]. Subsequently, the 2 µm square bins falling within the segmentation boundaries are aggregated to construct the transcriptional profile of each segmented unit. 2) **Refinement:** The initial segmentation output is fed into a probabilistic cell simulation using Proseg [9], which refines both the spatial boundaries and the transcriptional profiles of the segmented units. 3) **Downstream Processing and Export:** The refined units are exported into .loupe format for interactive visualization via the 10X Loupe Browser, as well as the .zarr open-standard format for large multidimensional arrays. Additionally, TENGU features a computational module to assist with cell annotation using user-selected, curated scRNA-seq models, such as CellTypist [13]. Any resulting cell type predictions are seamlessly integrated into the exported objects. Finally, TENGU includes a module to calculate spatially constrained cell-cell communication (CCC) using ligand-receptor interaction data from a user-specified database [19].

**Figure 1:**
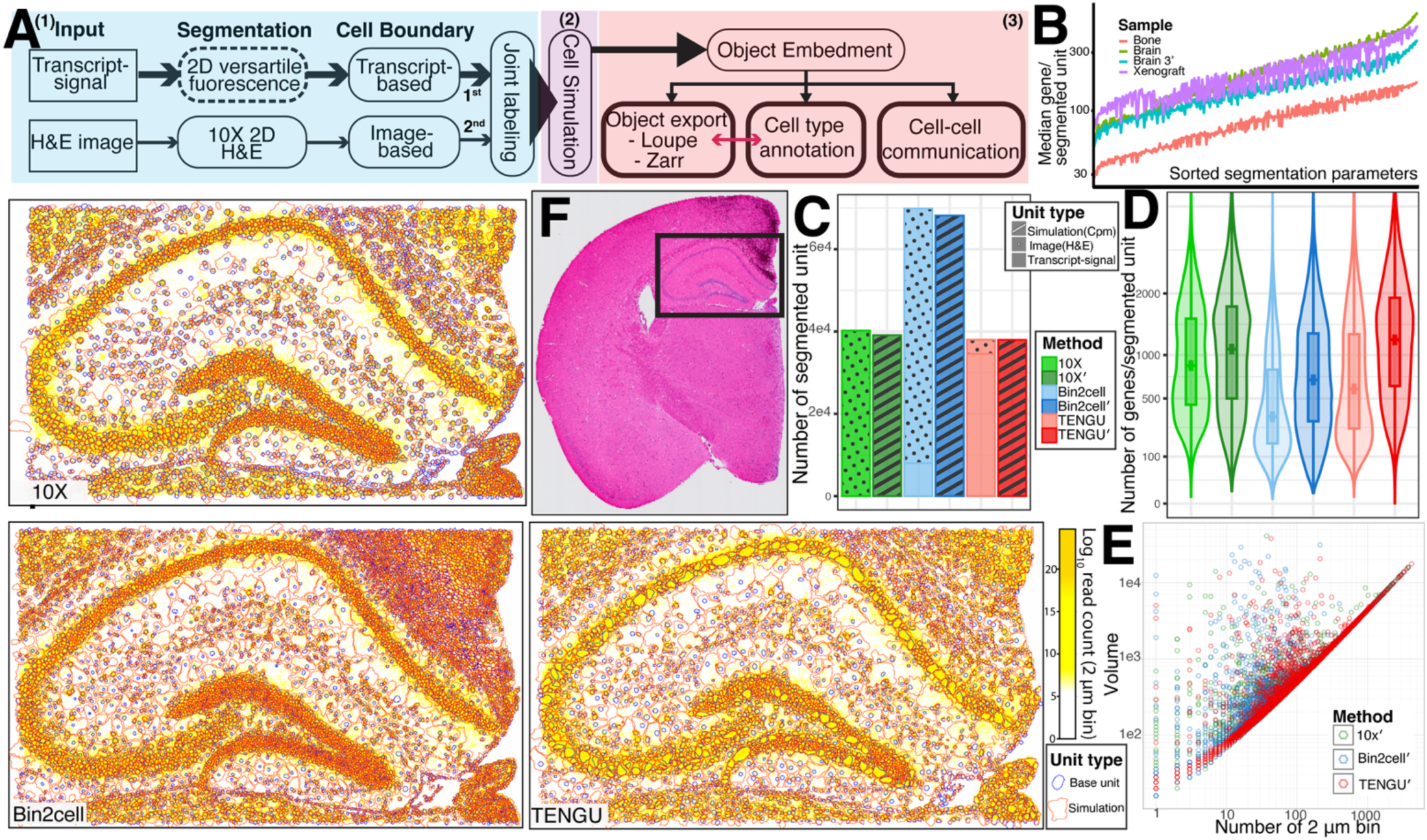
Overview of the TENGU workflow and comparative performance in high-resolution spatial transcriptomics. (A) Schematic overview of the TENGU pipeline. The workflow consists of three main stages: (1) an initial segmentation step that integrates boundaries derived from transcript-signal segmentation (using 2D versatile fluorescence with optimized parameters) and H&E image segmentation, (2) a cell simulation step to refine and aggregate boundaries and transcript profiles, and (3) downstream object embedding for export, cell type annotation, and cell-cell communication analysis.(B) Evaluation of segmentation parameters (several combinations) across diverse tissue samples (Bone, Brain, Brain 3’, and Xenograft), demonstrating the relationship between parameter tuning with objective function of median genes per segmented unit.(C) Bar plot comparing the total number of segmented units generated by the 10X, Bin2cell, and TENGU methods, both before and after the application of cell simulation (simulated outputs are denoted with a prime notation). Fill patterns indicate the source data driving the unit definition (Transcript-signal, H&E Image, or Simulation). (D) Violin-boxplot plots displaying the distribution of detected genes per segmented unit for each method. The prime notation (’) highlights the global increase in median gene detection following cell simulation. (E) Log-log scatter plot comparing the simulated cell volume against the number of constituents 2 µm bins for 10X’, Bin2cell’, and TENGU’. TENGU’ accurately maintains the theoretical minimum volume, indicated by the strict diagonal baseline, whereas alternative methods display upward scatter indicative of boundary overestimation. (F) H&E stained whole-tissue image of the mouse brain section used for the case study. The black rectangle delineates the highly cellular hippocampus region depicted in the adjacent spatial plots. (Left & Bottom) High-resolution spatial maps comparing segmentation outputs for 10x, Bin2cell, and TENGU within the hippocampus. Base segmented units (blue outlines) and post-simulation cell boundaries (orange outlines) are overlaid on a background heatmap representing the log read count per 2 µm bins.

### Comparative Analysis of Cell Segmentation and Simulation Approaches

Next, we performed comparative analysis of segmentation results on a brain tissue Visium HD dataset (probe based) derived from TENGU, Bin2cell, and 10X (See supplementary Fig S1 for workflow summary of Bin2cell and 10x). The Bin2cell and 10x methods rely on the 2D versatile models of H&E image and fluorescence provided as standard model from StarDist software package. Bin2cell uses segmentation results from H&E images as the primary and supplement with segmentation results from transcript-signal as secondary. 10X developed a custom H&E StarDist v6 model in-house derived from over 17,500 H&E image patches obtained from >150 H&E section images from a variety of tissues and preservation methods and used the model for segmentation. Subsequently, the 2 µm bins falling within the segmentation boundaries derived from the two methods were aggregated to construct the transcriptional profile of each segmented unit.

Bin2cell yielded the highest number of segmented units, identifying over 60,000, compared to TENGU and the 10x approach, which both yielded approximately 40,000 units (Figure 1C). Notably, 91% of TENGU’s segmented units were directly supported by underlying transcript signals, in contrast to only 12% for Bin2cell. Following cell simulation, we observed a minor reduction in the total number of segmented units for the 10x (1,083 units) and Bin2cell (1,683 units) methods, whereas TENGU was unaffected (Figure 1D). This reduction in 10x and Bin2cell was primarily driven by the elimination of false-positive segmentation artifacts (Supplementary Figure S2). Because the simulation algorithm expands the boundaries of segmented units by aggregating nearby, transcriptionally similar signals (Figure 1F), there was a global increase in the median number of genes per segmented unit compared to the original segmentations (Figure 1D).

Next, we compared the simulated cell volumes, derived from cell simulation, against the number of constituent 2 µm bins assigned to each segmented unit across the three methods (Figure 1E). In this log-log plot, a strict linear correlation at the lower boundary represents the theoretical minimum volume, where the simulated boundary is tightly restricted to the physical area of the assigned spatial bins. While this baseline establishes the theoretical minimum, we still observed some inflation of total volume across all three methods. This inflation occurs when simulated boundaries are expanded into empty intercellular spaces (sparse cell areas) and tissue edges, leading to a possible overestimation of cell size for a subset of segmented units. We then compared cell segmentation and simulation outputs derived from the three approaches. Focusing on the highly cellular regions of the hippocampus (Figure 1F) containing tightly packed as well as loosely organized anatomical structures. The basal transcript signal from 2 µm bins was visualized as a background density heatmap to serve as a ground truth for molecular location. For each analytical approach, the initial cell segmentation boundaries (blue) and the resulting simulated cell boundaries (red) were overlaid onto the transcript signal. Visual assessment of the cell layers, such as the dentate gyrus, reveals distinct differences in spatial resolution among the tools. While the 10x approach captures the general cellular organization, the simulated cell boundaries (red) occasionally deviate from the densest transcript regions. Conversely, the Bin2cell approach generates simulated boundaries that appear overly dense and generalized, struggling to maintain the distinct, empty intercellular spaces and potentially overestimating cell volume. In contrast, TENGU demonstrates high morphological fidelity. The simulated cell boundaries (red) generated by TENGU maintain a strict concordance with both the core cell boundaries (blue) and the underlying transcript density. Furthermore, by preventing the artificial merging of neighboring cells in tightly packed anatomical structures that cannot be discriminated by transcript signals alone, TENGU groups cells that share similar transcriptional profiles into a segmented unit and further denoise by the cell simulation.

### Cell Simulation Enhances Transcriptomic Signatures and Spatial Concordance

To evaluate the biological accuracy of the transcriptomic profiles generated by each segmentation approach, we performed automated cell type annotation using the CellTypist model and data from scRNA-seq of mouse brain [20] as well as a comprehensive brain scRNA-seq reference [21]. We specifically assessed the impact of our cell simulation algorithm by comparing the base segmentations against their simulated counterparts (denoted with a prime notation). An analysis of the cell type prediction confidence scores for 20 selected cell types (11 exhibiting a median score > 0.5 in at least one method, and 9 chosen based on their distinct topological structures within the MERFISH reference [21]) revealed that applying the cell simulation step consistently enhanced prediction confidence across all methods (Figure 2A). Notably, TENGU with cell simulation (TENGU’) achieved the highest overall median confidence assignments across a broad spectrum of identified cell types, markedly outperforming both its pre-simulation baseline (TENGU) and the simulated outputs of the alternative methods (10X’ and Bin2cell’).

**Figure 2:**
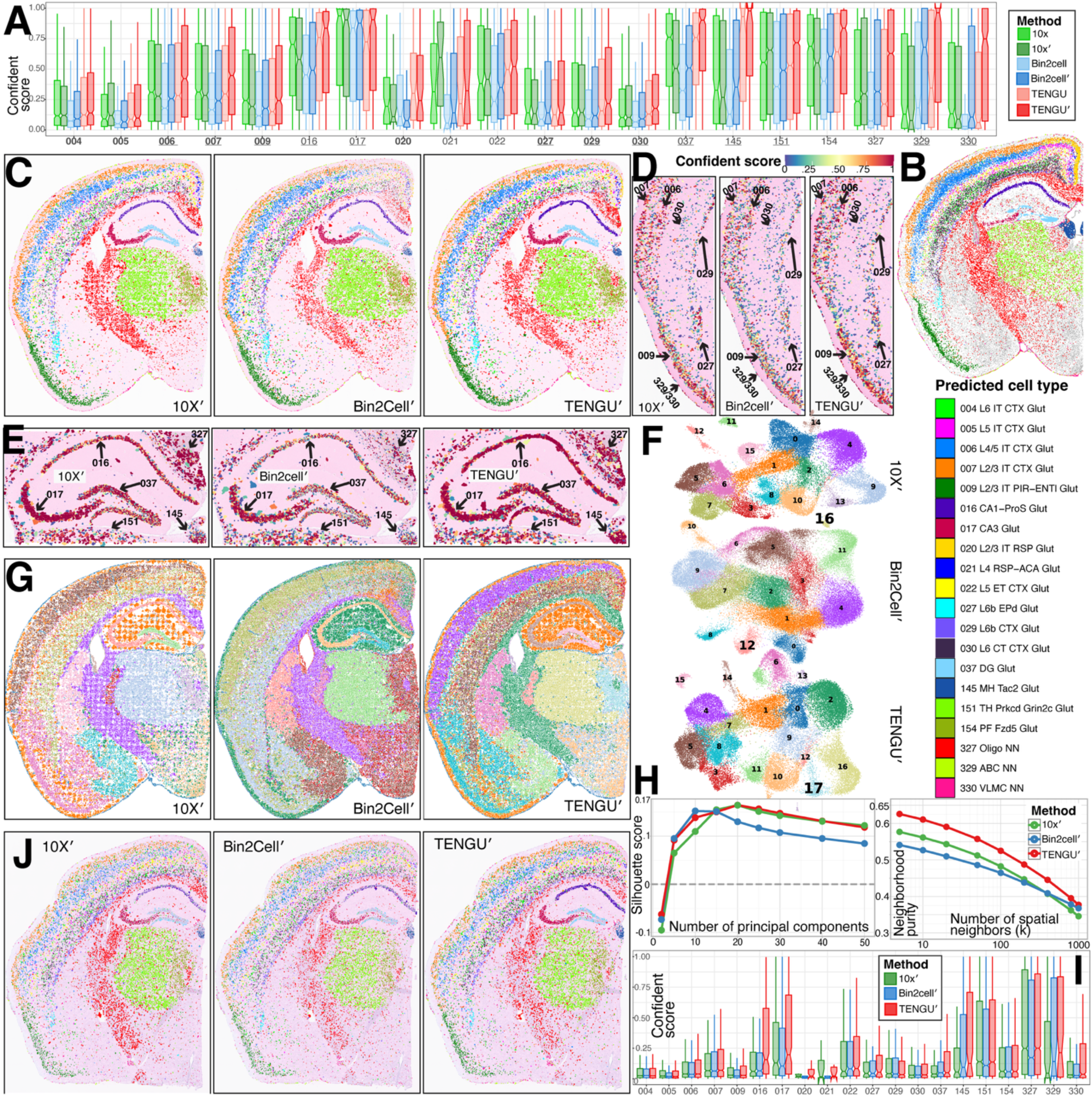
Cell simulation enhances transcriptomic signatures and spatial concordance. (A)Cell type prediction confidence scores derived from the scRNA-seq CellTypist model for 20 selected cell types, comparing base segmentations to their simulated counterparts (denoted with a prime, ’). TENGU’ achieves the highest overall median confidence assignments, outperforming its pre-simulation baseline and the simulated outputs of 10X’ and Bin2cell’. Underlined cell types are shown at higher magnification in panels D and E. (B) Matched MERFISH spatial transcriptomics dataset serving as a high-resolution, single-cell ground truth for anatomical architecture, with individual predicted cell types indicated by color. (C) Spatial mapping of predicted cell types for 10X’, Bin2cell’, and TENGU’, demonstrating overall spatial concordance with the MERFISH reference. (D) High-magnification spatial confidence heatmaps of cortical layers. Despite overall cell type prediction scores being relatively low, they preserve high spatial location concordance with the MERFISH reference. (E) Detailed spatial mapping of the hippocampal formation. TENGU’ accurately localizes specialized neuronal subtypes, notably CA3 (cluster 017) and Dentate Gyrus (DG) (cluster 037) glutamatergic neurons, with distinct spatial boundaries mirroring the MERFISH data. (F) UMAP embeddings of standard unsupervised clustering on the same parameter. TENGU’ delineates 17 distinct transcriptomic clusters, whereas 10X’ and Bin2cell’, yielding 16 and 12 clusters, respectively. (G) Spatial projection of the unsupervised clusters from (F) onto the physical tissue space. TENGU’ yields contiguous, biologically plausible spatial domains, minimizing the transcriptomic mixing and spatial scattering observed in 10X’ and Bin2cell’. (H) Quantitative assessment of transcriptomic distinctness and spatial coherence. Left: Silhouette score within the principal component (PC) space, with TENGU’ and 10X’ peaking optimally around 20 PCs (∼0.17). Right: Neighborhood purity score across a range of spatial neighbors (k), where TENGU’ retains a superior spatial signal-to-noise ratio. (I) Cell type prediction confidence scores derived from the 3’ poly-A chemistry dataset. (J) Spatial mapping corresponding to the 3’ poly-A chemistry dataset. While the cellular topology is biologically acceptable and maintains contiguous tissue organization, it exhibits lower spatial resolution compared to (C). Despite these dataset constraints, TENGU’ continues to outperform alternative methods.

To validate the spatial accuracy of the simulated profiles, we compared the mapped cell types against a matched MERFISH spatial transcriptomics dataset [21], which served as a high-resolution, single-cell ground truth (Figure 2B). Mapping the predicted cell types back to their spatial coordinates (Figure 2C) revealed that all three methods generally exhibited good spatial concordance with the established anatomical architecture delineated by the MERFISH reference, even in instances where their cell type prediction confidence scores were low. Closer inspection of specific anatomical regions of cortical layers (Figure 2D), we observed that despite yielding relatively low overall cell-type prediction scores, the spatial distributions of these populations remained highly concordant with the MERFISH ground truth. Furthermore, analysis of the hippocampal formation highlighted the enhanced spatial resolution uniquely provided by TENGU’ (Figure 2E). TENGU’ accurately localized specialized neuronal subtypes such as CA3 and Dentate Gyrus (DG) glutamatergic neurons (clusters 017 and 037, respectively) with distinct spatial boundaries that closely mirrored the MERFISH data. Furthermore, the spatial confidence heatmaps visually confirmed that while the overall topologies were captured across methods, the cell type assignments of TENGU’ within tightly packed anatomical regions possessed markedly higher prediction scores compare to the dispersed, lower-confidence assignments produced by 10X’ and Bin2cell’.

To complement the automated annotations and evaluate the clustering capacity of each method, we performed standard unsupervised clustering using identical calculation parameters and visualized the overall transcriptomic landscape via UMAP embeddings (Figure 2F). Under a fixed clustering resolution, we observed clear differences in the number of distinct populations successfully recovered by each pipeline. Specifically, TENGU’ delineated 17 distinct transcriptomic clusters across the UMAP embedding, whereas the noisier transcriptomic profiles of 10x’ and Bin2cell’ resulted in the artificial merging of closely related states, yielding only 16 and 12 clusters, respectively. To quantitatively assess this transcriptomic distinctness, we calculated the Silhouette score within the high-dimensional principal component (PC) space based on the clustering results (Figure 2H, left). TENGU’ and 10x’ achieved the highest overall Silhouette scores, peaking optimally around 20 principal components and reaching values (approximately 0.17) comparable to those previously reported for complex scRNA-seq data [22, 23], markedly outperforming Bin2cell’. Together, this increased cluster recovery and high-dimensional separation indicate that TENGU’ and 10X’ better preserve the distinct molecular signatures required for robust downstream clustering compared to Bin2cell’. When projecting these unsupervised clusters back onto the tissue space, TENGU’ yielded contiguous, biologically plausible spatial domains that reflected known mouse brain anatomy [14, 21, 24, 25] (Figure 2G). In contrast, the spatial clusters generated by 10X’ and Bin2cell’ exhibited noticeable transcriptomic mixing and spatial scattering. To quantify this observation, we calculated the Neighborhood Purity Score across a range of spatial neighbors (k) (Figure 2H, right). TENGU’ consistently outperformed both Bin2cell’ and 10X’, maintaining a significantly higher purity score (starting above 0.60 at k=5). While all methods showed a decrease in purity as the neighborhood size increased toward k=1000, TENGU’ retained a superior signal-to-noise ratio throughout the entire scale. Together, these results indicate that TENGU’ effectively aggregates relevant local signals while minimizing transcriptomic bleed-through between neighboring cells, preserving the distinct molecular signatures required for robust clustering and highly confident cell-type identification.

To further evaluate performance across different assay chemistries, we examined the clustering and spatial mapping results derived from the 3’ poly-A chemistry of mouse brain tissue (Figures 2J and 2I). When compared to the initial spatial and transcriptomic profiles (Figure 2B), this sample yielded noticeably lower overall quantitative scores (compare to Figure 2A). However, despite these reduced metrics, the resulting cellular topology remained biologically acceptable and maintained contiguous tissue organization. While this spatial structure was functional, it did not quite achieve the robust topological clarity and high-resolution spatial definition demonstrated in Figure 2C. Crucially, even within the constraints of this dataset, TENGU’ continued to yield the best results, successfully outperforming the other methods.

### Spatial Transcriptomic Analysis of the Bone Microenvironment

To evaluate the performance of TENGU’ in tissues with high technical barriers, we applied our pipeline to decalcified mouse bone, a notoriously difficult tissue for spatial transcriptomics due to its dense, mineralized extracellular matrix and low cellularity. The sample used for this study was a femur from a murine model of *Staphylococcus aureus* osteomyelitis, which include cortical bone, bone marrow, and an abscess involving the marrow, cortical bone, and extending outside the bone (Figure 3A) [26]. TENGU’ successfully segmented the transcriptomic data, maintaining a robust number of segmented units while substantially increasing the median number of genes detected per unit compared to the 10X’ pipeline (Figure 3B). The segmentation and cell simulation results were exported as .loupe file and visualized in Loupe Browser software. This enriched transcriptomic recovery translated directly into superior clustering resolution using identical calculation parameters, mapping the unsupervised clusters back onto the spatial coordinates revealed that TENGU’ delineated 25 distinct cellular populations, whereas the noisier 10X’ pipeline resolved only 21 (Figure 3A).

**Figure 3:**
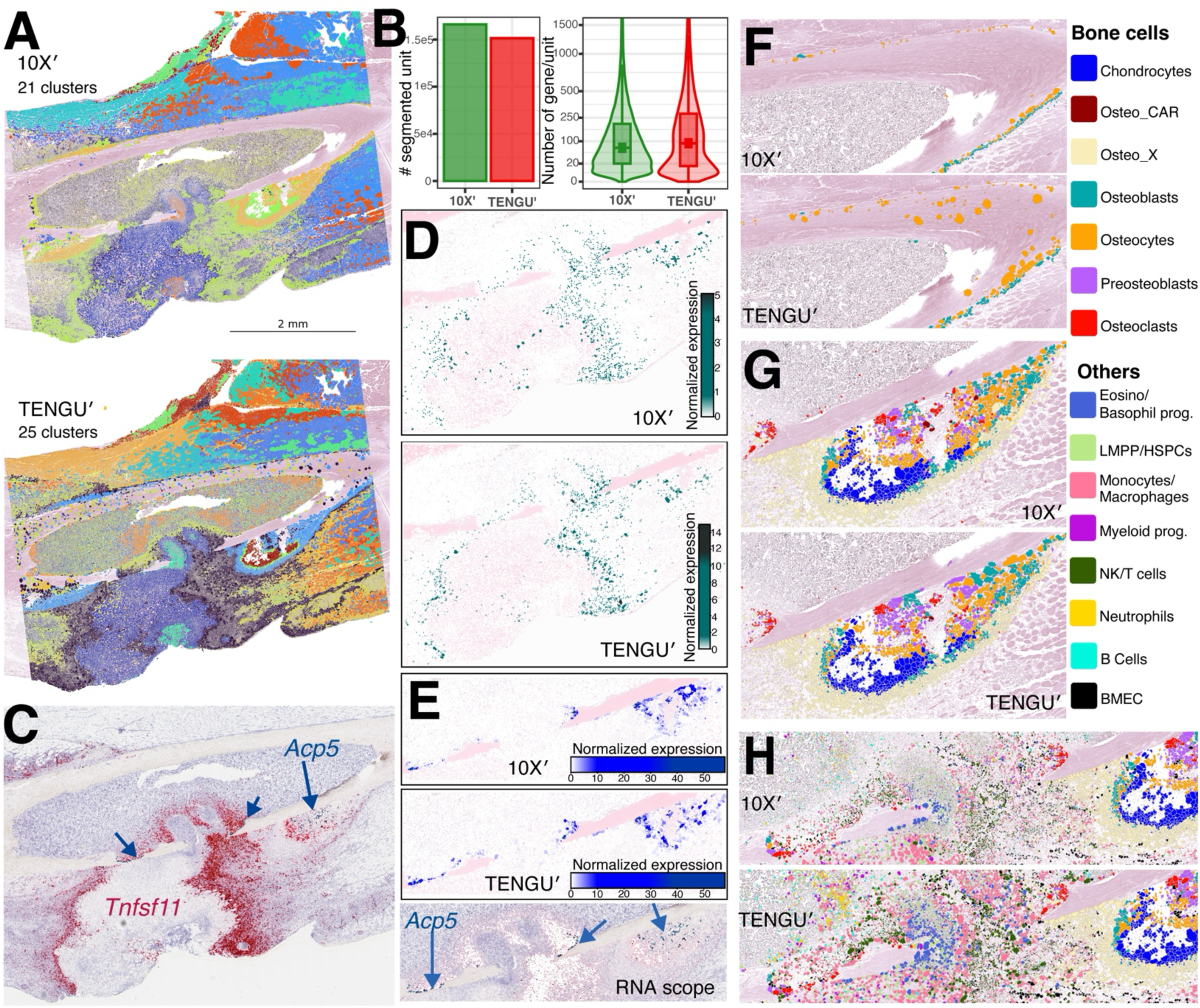
Spatial transcriptomic analysis and high-resolution mapping of the bone microenvironment. The results of segmentation and simulation were export as .loupe file and visualize in Loupe Browser with different arbitrary subset selection of predicted cell types. (A) Spatial projection of unsupervised clustering results on decalcified mouse femur from a murine model of *Staphylococcus aureus* osteomyelitis. TENGU’ delineates 25 distinct cellular populations compared to the 21 populations resolved by the standard 10X’ pipeline using identical clustering parameters. (B) Quantification of segmentation performance. Bar chart (left) showing the total number of segmented units recovered, and violin-box plot (right) demonstrating the substantial increase in the median number of genes detected per unit by TENGU’ versus 10X’. (C) RNAscope, in situ hybridization establishing the spatial ground truth for key markers *Tnfsf11* (RANKL, red) and *Acp5* (TRAP, blue). (D) Computationally derived normalized spatial expression of *Tnfsf11*, highlighting sharper biological boundaries and stronger signal localization using TENGU’ compared to 10X’. (E) Normalized spatial expression of *Acp5* across both computational pipelines, accurately mirroring the RNAscope validation. (F) Magnified spatial cell-type mapping of the cortical bone using a curated CellTypist model. TENGU’ successfully captures transcriptomic signatures from mature osteocytes embedded deep within the mineralized matrix, overcoming the severe signal dropout observed in the H&E-reliant 10X’ method. (G) High-resolution mapping of the delicate periosteum high bone remodeling niche, successfully preserving and resolving complex layers of chondrocytes, osteo-CAR cells, preosteoblasts, osteoblasts, osteocytes, osteoclasts surrounded by Osteo-X cells. (H) Magnified view of an active bone break and highly heterogeneous marrow cavity. TENGU’ accurately delineates the spatial co-localization and osteoimmune interactions between bone-resorbing osteoclasts and diverse hematopoietic/immune populations, including LMPP/HSPCs, monocytes/macrophages, neutrophils, and B cells, NK/T cells, progenitors (prog.) and bone marrow endothelial cells (BMEC).

### In Situ Validation and Detection of Matrix-Embedded Osteocytes

The bone osteomyelitis sample contains intense, localized inflammation and infection-induced bone resorption [26]. We compared the computationally derived expression profiles against RNAscope in situ hybridization of an adjacent section (Figure 3C). TENGU’ provided highly accurate, localized expression mapping of key markers driving this pathology. For instance, the normalized spatial expression of *Tnfsf11* (RANKL), a critical cytokine upregulated during osteomyelitis to stimulate osteoclastogenesis, was better captured by TENGU’. Our pipeline delineated sharper biological boundaries and a stronger expression signal within the active remodeling lesions than the signal produced by 10X’ (Figure 3D). Furthermore, both TENGU’ and 10X’ accurately mapped expression of *Acp5*, a marker of osteoclasts, highlighting the populations of active, bone-resorbing osteoclasts and closely mirroring the RNAscope ground truth (Figure 3E).

Next, to annotate the cell types of the segmented units for further high-resolution analysis, we constructed a CellTypist model of bone-resident cells derived from a curated dataset from our previous study [16]. Importantly, the magnified view of the cortical bone highlights a major technical advantage of TENGU’, namely the successful detection of osteocytes embedded deep within the mineralized tissue (Figure 3F). Capturing transcriptomic signatures from mature osteocytes is an acknowledged challenge in spatial biology due to poor probe penetration and low RNA yield [27]. TENGU’ successfully overcomes this barrier, rendering transcriptomic signals in regions where the 10x’ method, which relies solely on H&E images for segmentation, suffers from severe dropout.

### Reactive Bone Formation and Osteoimmune Interactions

We next focused on the effect of infection on the bone periosteum known alternatively as reactive bone formation or periosteal reaction [28]. Little is known about the mechanisms that drive formation of this structure, which occurs on the periosteal surface at the edges of the abscess. TENGU’ and 10X’ successfully preserved and mapped the complex architecture of this structure, distinctly resolving layers of chondrocytes, Osteo-CAR, pre-osteoblasts, osteoblasts, osteocytes, and osteoclasts, all surrounded by Osteo-X cells (Figure 3G). Further, a focus on the area in which the level of bone destruction is highest, TENGU’ was able to map the highly heterogeneous marrow cavity (Figure 3H). Here, TENGU’ clearly delineated the spatial co-localization of bone-resorbing osteoclasts alongside a rich diversity of hematopoietic and immune populations, including monocytes/macrophages, neutrophils, B cells, and lymphoid progenitors (LMPP/HSPCs).

### Spatial Cell-Cell Communication Analysis in the Bone Microenvironment

Having established that TENGU’ provides superior transcriptomic distinctness and spatial resolution within the complex, mineralized bone microenvironment (Figure 3), we next sought to leverage this high-fidelity segmentation to map spatially aware cell-cell communication (CCC) networks. For this analysis, we utilized a curated ligand-receptor database from a previous study [29], comprising 11,395 potential interactions. We first calculated comprehensive ligand-receptor interaction scores across all segmented units, factoring in both expression levels and spatial proximity based on the empirical formulation.

To anchor our proximity metrics, we firstly determined the nearest neighbor distance across all segmented cells within the tissue space (Figure 4A). This established the baseline physical architecture of the cellular network, revealing a median cell-to-cell distance of 46 μm. Because paracrine signaling relies on biological diffusion gradients, we iteratively calculated CCC scores across expanding neighborhood radii (ranging from the baseline of 46 μm up to 1,474 μm) to evaluate how spatial constraints affect the detection of functional ligand-receptor pairs (Figure 4B). As expected, we observed a robust increase in both the frequency and diversity of detected interactions as the radius expanded. However, this diversity plateaued at approximately 8,400 interactions, extending the radius beyond ∼737 μm yielded no significant gain in novel ligand-receptor types. This suggests that a 737 μm threshold effectively captures the maximum functional biological signaling distance within this specific tissue context.

**Figure 4:**
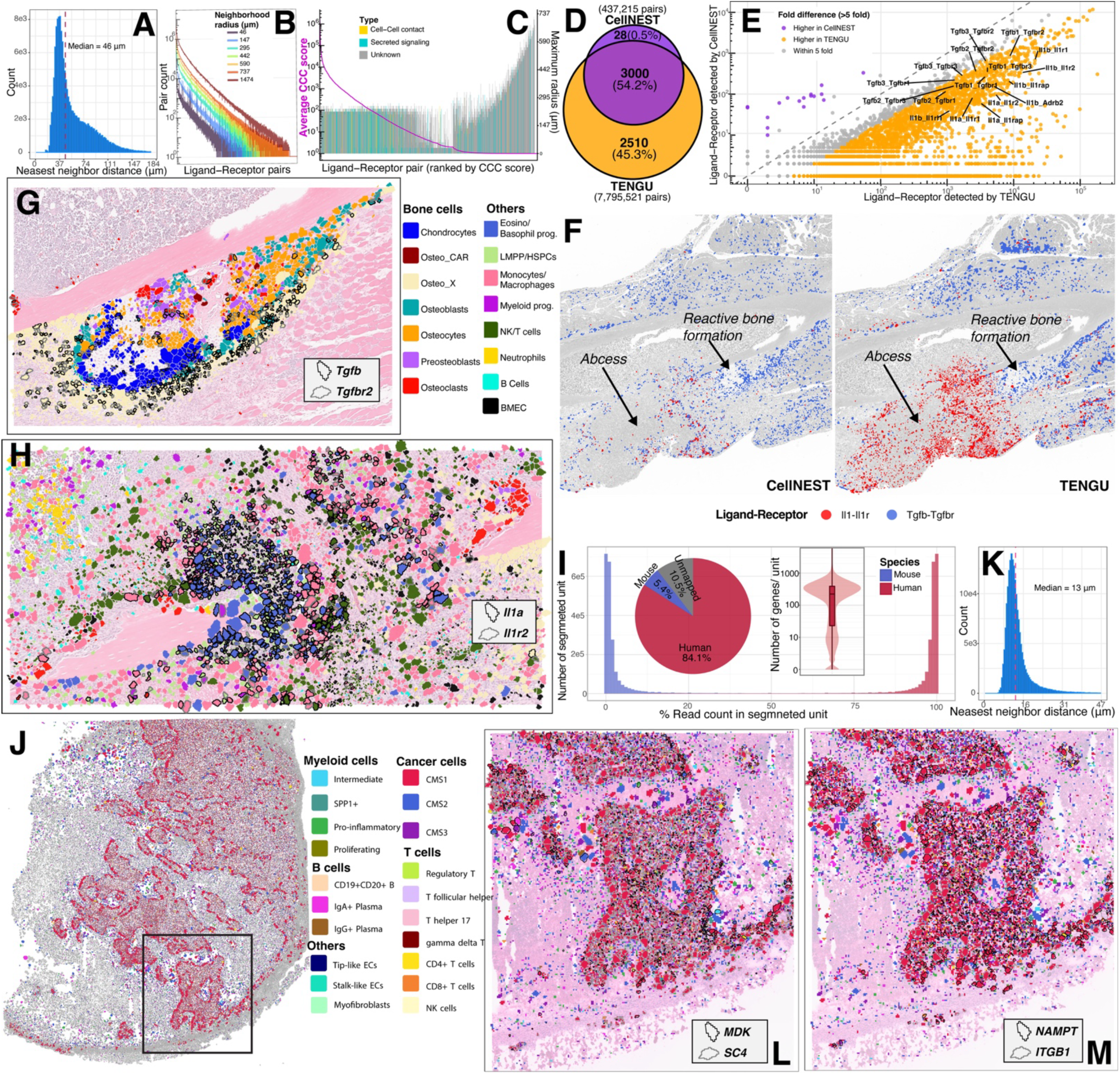
Spatial cell-cell communication (CCC) networks in the pathological bone microenvironment and dense tumor xenografts. (A) Distribution of the nearest neighbor distances across all segmented units within the tissue space. The red dashed line indicates the median physical distance of 46 μm, establishing the baseline cellular architecture. (B) Quantification of detected ligand-receptor interaction frequency and diversity evaluated iteratively across expanding neighborhood diffusion radii (ranging from 3.1 to 100 μm). The total number of unique functional interactions plateaus at a radius of approximately 50 μm. (C) Evaluation of spatial signaling modalities. The plot displays the overall CCC interaction score (magenta line, left y-axis) alongside the maximum functional spatial radius (bar height, right y-axis) for ranked ligand-receptor pairs, categorized by signaling mechanism (Cell-Cell Contact, Secreted Signaling, or Unknown). (D) Venn diagram comparing the distinct ligand-receptor pairs detected by CellNEST and TENGU. TENGU identifies a substantial proportion of unique interactions (45.3%) compared to CellNEST (0.5%). (E) Scatter plot showing the fold difference in ligand-receptor pair detection between the two tools. Pairs detected at >5-fold higher frequency in TENGU are highlighted in orange, including key interleukin-1 interactions (e.g., Il1b_Il1r1, Il1a_Il1r1) and TGF-beta interactions (e.g., tgfb1, tgfb2, tgfb3, tgfbr1, tgfbr2, tgfbr3) , while pairs higher in CellNEST are shown in purple. (F) Spatial mapping of Il1-Il1r (red) and Tgfb-Tgfbr (blue) signaling interactions on bone tissue sections. Both tools map TGF-beta$signaling to areas of reactive bone formation. However, TENGU distinctly captures the dense, localized concentration of IL-1 signaling encapsulating the abscess, which is detected at a much lower density in the CellNEST analysis. This spatial distribution accurately reflects established models of bone infection, where IL-1-driven abscess encapsulation directly triggers the adjacent reactive bone remodeling captured by the TGF-$\beta$ markers. This spatial distribution accurately reflects established models of bone infection, where IL-1-driven abscess encapsulation directly triggers the adjacent reactive bone remodeling captured by the TGF-beta markers. (G) High-resolution spatial mapping of the osteogenic Tgfb (black boundaries) and Tgfbr2 (gray boundaries) signaling axis within the dense periosteal remodeling zone. TENGU’ successfully resolves the localized sender and receiver populations, highlighting intense crosstalk among resident bone cells. Cell type annotations are provided in the accompanying legend. (H) Spatial visualization of the pro-inflammatory Il1a (black boundaries) and Il1r2 (gray boundaries) signaling network within the highly infiltrated marrow cavity characteristic of the *S. aureus* osteomyelitis model. The mapping captures complex, long-range immune-stromal and immune-immune interactions driven by recruited myeloid populations. (I) Species-specific transcriptomic recovery in a human colorectal cancer xenograft model. The pie chart displays the global read mapping distribution, revealing dominant human cancer cell activity (84.1%). The histogram demonstrates the highly skewed, bimodal distribution of human reads per segmented unit. Applying a >95% human read cutoff to successfully isolate the tumor compartment, the resulting distribution of detected genes per segmented unit is illustrated in the inset violin-boxplot. (G) Spatial cell-type annotation of the filtered human tumor microenvironment using a human colorectal cancer CellTypist model. TENGU’ resolves diverse cancer cell states and human immune/stromal populations. The square box indicate the region of interest. (K) Nearest neighbor distance distribution of the segmented units within the xenograft. The median distance of 13 µm (red dashed line) highlights the extreme cellular density of the tumor compared to the bone tissue. (L & M) High-resolution CCC spatial mapping within the dense tumor mass. TENGU’ successfully isolates localized, species-specific signaling hubs without spatial bleed-through, specifically highlighting the pro-tumorigenic MDK-SDC4 axis (L) and the NAMPT-ITGB1 interaction network (M).

### Distinguishing Spatial Signaling Modalities Based on Functional Radius

To further characterize the biophysical nature of the identified communication networks, we evaluated the relationship between the interaction strength (CCC score) and the maximum functional spatial radius required to achieve that interaction for each specific ligand-receptor pair (Figure 4C). By categorizing these interactions by their established signaling mechanisms based on the database [29], our spatial framework successfully differentiated between localized, contact-dependent interactions and broader, diffusion-based networks. As anticipated, known pathways classified as using direct cell-cell contact generally operated within tighter spatial constraints, correlating with shorter maximum functional radii. Conversely, secreted signaling pathways demonstrated a significantly broader range of effective communication distances across the tissue architecture. Notably, by simultaneously tracking the interaction score and spatial reach, TENGU’ provides a data-driven methodology to infer the biophysical limits of uncharacterized or “unknown” ligand-receptor pairs.

### Benchmarking TENGU against CellNEST in spatial-aware CCC analysis

To benchmark the CCC detection capabilities of TENGU against existing state-of-the-art method, we compared its performance with CellNEST [18] on the bone tissue based on the same ligand-receptor (LR) dataset. CellNEST was selected as the benchmark due to it represents a major advancement over earlier spatial-aware CCC tools; by utilizing deep learning attention mechanisms. However, TENGU demonstrated vastly superior sensitivity in our biological model (Figure 4D), detecting 7,795,521 total LR pairs compared to 437,215 pairs detected by CellNEST. While both tools successfully identified a shared core of 3,000 distinct LR interactions (54.2%), TENGU uniquely identified 2,510 additional interactions (45.3%), whereas CellNEST only yielded 28 unique interactions (0.5%). Quantitative comparison of detection frequencies revealed a strong skew toward TENGU, with a vast majority of identified LR pairs showing greater than 5-fold higher detection in TENGU (Figure 4E). To assess the biological accuracy of these findings, we spatially mapped key inflammatory and bone remodeling signaling pathways onto the tissue (Figure 4F). Both TENGU and CellNEST successfully localized TGF-beta signaling (Tgfb-Tgfbr) to regions of reactive bone formation. However, CellNEST detected substantially less IL-1 signaling (Il1-Il1r) within the tissue. In contrast, TENGU robustly identified dense, highly localized IL-1 signaling directly surrounding the abscess. The accurate spatial capture of this IL-1 signaling by TENGU is biologically critical; localized IL-1 beta and IL-1R activation is a well-established driver of abscess encapsulation in infectious osteomyelitis [30, 31]. Furthermore, this concentrated IL-1 signaling stimulates local osteoclast-driven bone resorption, which mechanically triggers the compensatory reactive bone formation observed in the adjacent tissue. TENGU thus accurately resolves the complete pro-inflammatory and structural remodeling microenvironment that is only sparsely captured by CellNEST.

### Mapping of Osteogenic and Inflammatory Signaling Hubs

To demonstrate the biological utility of our basic spatially aware CCC framework, we visualized targeted ligand-receptor interactions within distinct pathological and remodeling niches. First, we focused on the reactive bone formation area of the osteomyelitis sample (Figure 4G). Here, we mapped the spatial distribution of the *Tgfb* (transforming growth factor-beta) and *Tgfbr2* (TGF-beta receptor 2) signaling axis, a fundamental pathway for osteoblastogenesis and bone repair [32]. TENGU’ successfully resolved a tight spatial signaling hub, clearly delineating the specific sender (ligand-expressing) and receiver (receptor-expressing) populations. The interaction was highly localized to the Osteo-X population of cells [16], forming the outer edge of the reactive bone formation structure (Figure 4G). These results suggest that Tgfb autocrine or paracrine signaling is important for the function of Osteo-X cell during the process of reactive bone formation and highlight the ability of TENGU to detect a specific CCC within a complex and highly active environment.

Next, we examined the highly heterogeneous marrow cavity, which is characterized by intense immune infiltration driven by the *S. aureus* osteomyelitis model (Figure 4H). In this inflammatory niche, we evaluated the spatial interaction between the potent pro-inflammatory cytokine *Il1a* (Interleukin-1 alpha) and its receptor *Il1r2*. Unlike the structurally confined TGF-β signaling in the periosteum, the *Il1a*–*Il1r2* axis revealed a distinct spatial architecture driven by immune-stromal and immune-immune crosstalk. TENGU’ mapped these interactions across the heavily infiltrated marrow space, capturing the complex, localized signaling networks between recruited myeloid populations (e.g., neutrophils and monocytes/macrophages) and the surrounding microenvironment. Together, these spatial visualizations demonstrated that TENGU’ effectively decodes the functional micro-anatomy of both tissue repair and active pathological inflammation.

### Species-Specific Spatial Resolution in a Dense Tumor Microenvironment

To further demonstrate the versatility of TENGU’ across diverse tissue architectures and sequencing chemistries, we applied our pipeline to a human colorectal cancer (CRC) xenograft model implanted in the mouse colon. This tissue was processed using 3’ poly(A) capture-based chemistry, which inherently captures transcripts from both organisms simultaneously. Evaluation of the global mapping metrics revealed that the human cancer cells were vastly more transcriptionally active than the murine host tissue, accounting for 84.1% of the total mapped reads. This overall dominance resulted in a highly skewed distribution, with individual segmented units overwhelmingly populated by human, rather than murine, transcripts (Figure 4I, see Supplementary Fig S3 for a spatial plot). To eliminate cross-species transcriptomic bleed-through and focus strictly on the intratumoral signaling dynamics, we established a stringent species-specificity threshold, retaining only segmented units containing >95% human reads. The number of genes detected per segmented unit is illustrated in the accompanying violin plot.

Using a CellTypist model trained on human CRC single-cell data [15], we successfully mapped the complex spatial heterogeneity of the human tumor compartment. TENGU’ identified distinct malignant epithelial states alongside infiltrating human immune and stromal populations, capturing the compartmentalized architecture of the tumor microenvironment (TME) (Figure 4J). Notably, calculating the nearest neighbor distance within this tumor highlighted a drastically different physical constraint compared to the bone microenvironment. The median cell-to-cell distance in the xenograft was just 13 µm (Figure 4K), reflecting the hyper-proliferative, compacted nature of a solid epithelial tumor.

### Mapping Pro-Tumorigenic and Metabolic Signaling Hubs

Despite this high cellular density, TENGU’ successfully resolved spatial cell-cell communication networks. Zooming into the active tumor regions, we mapped the localized spatial distribution of the MDK (Midkine) and SDC4 (Syndecan-4) signaling axis (Figure 4L). Midkine is a potent heparin-binding growth factor, and its interaction with the SDC4 proteoglycan receptor is a critical driver of CRC proliferation, angiogenesis, and focal adhesion remodeling [33]. Crucially, TENGU’ delineated spatially distinct sender and receiver populations, MDK ligand expression (black boundaries) was highly concentrated within the dense islands of malignant epithelial cells, while SDC4 receptor expression (gray boundaries) mapped predominantly to the adjacent stromal and endothelial margins. This clear spatial segregation visualizes a signaling axis driving outward tumor expansion. Similarly, we resolved the distinct spatial architecture of the NAMPT (Nicotinamide phosphoribosyltransferase) and ITGB1 (Integrin beta-1) interaction network (Figure 4M). Extracellular NAMPT signaling through integrin receptors serves as a crucial metabolic and inflammatory bridge in the TME, promoting cancer cell survival under metabolic stress and driving fibrotic stromal reprogramming [34]. Rather than ubiquitous co-localization, TENGU’ highlighted distinct pockets of NAMPT-expressing cells (senders) signaling across the tissue space to structurally widespread ITGB1-expressing stromal cells (receivers).

### Application of TENGU to High-Density Human Breast Cancer Tissue over Atera platform

To further evaluate the versatility of TENGU across emerging spatial transcriptomics platforms, we applied the pipeline to a human breast cancer tissue section profiled with the Atera platform. We focused our high-resolution analysis on a 2.8 mm × 4.65 mm subset (blue rectangular Figure 5A) contains both breast cancer cell type of ductal carcinoma in situ (DCIS) cells and invasive cells based on the transcriptional profile of 11q13-resident genes [35]. Consistent with our prior benchmarks, TENGU achieved robust transcriptomic recovery, yielding a median of 1,841 genes per segmented unit compared to 1,496 genes per unit for the standard 10X pipeline (Figure 5C). Cell-type annotation resolved 17 distinct epithelial, stromal, and immune populations within the breast tumor microenvironment. These included multiple 11q13 high-grade DCIS states (general, mitotic, and G1/S tumor cells), basal-like structured DCIS, luminal-like amorphous DCIS, apocrine cells, and myoepithelial cells, alongside high- and low-grade DCIS-associated cancer-associated fibroblasts (CAFs), CXCL14+ fibroblasts, endothelial cells, pericytes, and infiltrating immune populations (T lymphocytes, macrophages, myeloid cells, mast cells, and plasma/mast cell mixtures). infiltrating immune populations (T lymphocytes, macrophages, myeloid cells, mast cells, and plasma/mast cell mixtures).

**Figure 5.**
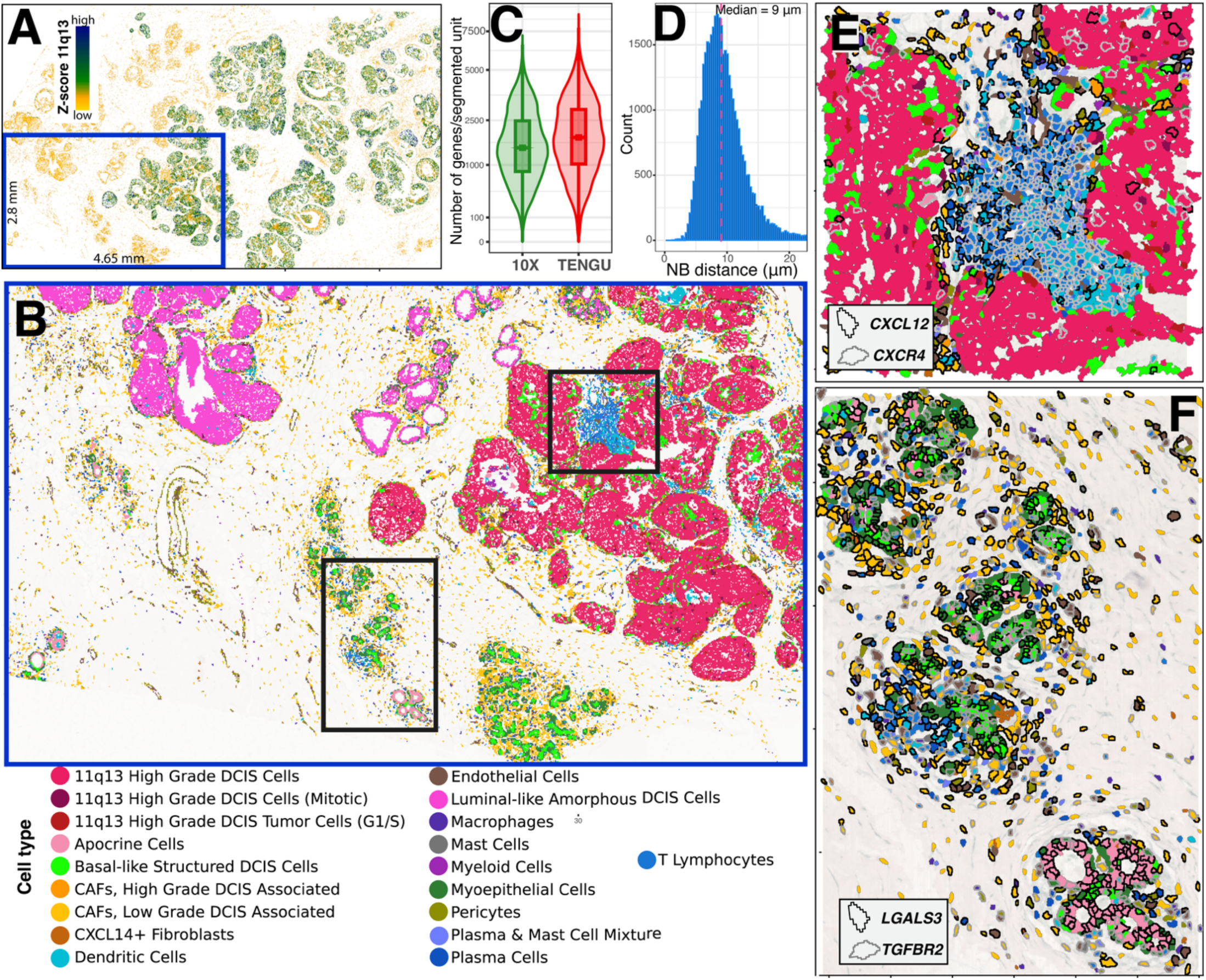
Spatial transcriptomic analysis and cell-cell communication mapping of high-density human breast cancer tissue profiled with Atera platform. (A) Spatial mapping of the 11q13 Z-score (low to high) based on transcriptional levels of genes within the locus, delineating the anatomical boundaries of high-grade ductal carcinoma in situ (DCIS) lesions. (B) High-resolution spatial cell-type annotation of a 2.8 mm × 4.65 mm tissue subset, resolving 17 distinct epithelial, stromal, and immune populations within the breast tumor microenvironment, color-coded as indicated in the accompanying legend. (C) Distribution of the number of detected genes per segmented unit comparing the standard 10X pipeline and TENGU, demonstrating enhanced transcriptomic recovery by TENGU (median of 1,841 vs. 1,496 genes per unit). (D) Distribution of nearest-neighbor (NB) distances across all segmented units within the tissue subset; the median physical distance of 9 µm highlights the high cellular density of the breast cancer microenvironment. (E) High-resolution spatial cell-cell communication (CCC) mapping of the CXCL12 (black boundaries) and CXCR4 (gray boundaries) chemokine signaling axis within the first region of interest (ROI), resolving CXCL12-expressing stromal fibroblasts and interface cells signaling to CXCR4-expressing immune infiltrates positioned between dense high-grade DCIS clusters. (F) Spatial visualization of the LGALS3 (black boundaries) and TGBR2 (gray boundaries) signaling network within the second ROI, capturing localized paracrine interactions between structured ductal/lobular epithelial compartments and adjacent stromal and immune populations.

Quantification of the local cellular architecture revealed a median nearest-neighbor distance of just 9 µm (Figure 5D), representing an even more densely compacted microenvironment than both the decalcified bone (46 µm) and the colorectal cancer xenograft (13 µm). Despite this high cellular density, TENGU’s spatially aware CCC module successfully resolved localized signaling networks across two distinct regions of interest (ROIs). In the first ROI, which captured dense islands of high-grade DCIS surrounding a central immune-infiltrated stromal corridor, we mapped the CXCL12– CXCR4 chemokine signaling axis (Figure 5E). In breast cancer, fibroblast-derived CXCL12 signaling through CXCR4 is a well-established driver of stromal desmoplasia, immune cell recruitment, and cytotoxic T-lymphocyte exclusion from malignant nests [36]. TENGU clearly segregated these sender and receiver compartments: CXCL12 ligand expression (black boundaries) was predominantly localized to stromal fibroblasts and cells along the tumor–stroma interface, whereas CXCR4 receptor expression (gray boundaries) was densely concentrated within the central infiltrate of immune cells positioned between the malignant DCIS clusters (Figure 5E). In the second ROI, which encompassed organized ductal and lobular structures embedded within the surrounding stroma, we evaluated the LGALS3-TGFBR2 signaling axis (Figure 5F). LGALS3 (encoding Galectin-3) is a multifunctional β-galactoside-binding lectin secreted by both mammary epithelial and infiltrating cells that promotes tumor cell survival, extracellular matrix remodeling, and local immunosuppression by modulating macrophage polarization and T-cell activity within the tumor niche [37]. Here, LGALS3 expression (black boundaries) was prominently distributed across the structured epithelial/ductal compartments and adjacent infiltrating stromal and immune cells, signaling locally to neighboring [Receptor]-expressing populations (gray boundaries) concentrated along the ductal margins and surrounding microenvironment (Figure 5F). Together, these results demonstrate that TENGU generalizes effectively to the Atera data, preserving boundary fidelity and resolving spatially distinct paracrine signaling hubs even within highly compacted breast cancer tissues.

## Discussion

The rapid adoption of high-resolution spatial transcriptomics, particularly the Visium HD and Atera platform, has created a need for robust computational frameworks capable of processing massive datasets without sacrificing biological fidelity. In this study, we developed TENGU, an end-to-end bioinformatic pipeline that successfully navigates the complex interplay between image registration, transcriptomic binning, and spatial interaction mapping. By demonstrating TENGU’s efficacy across highly diverse architectures, ranging from organized murine brain structures to sparse mineralized bone and highly compacted human tumor xenografts and human breast cancer, we highlight its adaptability and superior performance compared to existing standard pipelines.

A fundamental and persistent challenge in spatial transcriptomics is achieving true single-cell resolution. Standard tissue sections prepared for the Visium HD and Atera platform are typically 5–10 µm thick, meaning they often capture only a cross-sectional fragment of a cell rather than its entirety. Furthermore, the physical angle of tissue sectioning heavily influences how morphological boundaries are projected onto the 2D capture area. Consequently, a single computationally defined “segmented unit” may inadvertently encompass overlapping fragments from multiple distinct physical cells. However, TENGU applies a transcriptomic-driven cell simulation step using the robust framework Proseg [9] to minimize the transcriptomic mixing of different cell types in the segmented units by iteratively refining boundaries based on localized gene expression probabilities. Crucially, even in instances where a segmented unit physically merges multiple cells due to extreme tissue density, the simulation ensures that the aggregated spatial bins share highly correlated expression profiles. Therefore, the segmented unit is likely to represent a biologically homogeneous cell type. This mechanism preserves the transcriptomic distinctness necessary for accurate, high-resolution downstream clustering and improves confidence in cell-type predictions using scRNA-seq based models.

A common metric used to assess spatial transcriptomic data is the total number of transcripts detected per cell. However, achieving a transcriptomic yield comparable to standard single-cell RNA sequencing should not be considered a universal baseline, as transcript recovery is highly tissue-dependent. For instance, while soft tissues like brain may yield robust transcriptomic profiles approaching single-cell levels, tissues such as decalcified bone inherently suffer from much lower RNA yields. This reduction in the median number of transcripts detected per segmented unit is an unavoidable consequence of the harsh chemical processing required for decalcification and the physical barrier of the mineralized extracellular matrix. Consequently, while computational tools can maximize data recovery, further optimization of experimental and tissue preparation protocols remains necessary for these challenging microenvironments.

Beyond defining cellular identity, a primary goal of spatial biology is uncovering cell-cell communication within microenvironments. TENGU provides a streamlined, a basic module for spatially constrained cell-cell communication (CCC). We recognize that the bioinformatic landscape for CCC is rapidly expanding [38] This growth has resulted in the continuous emergence of highly specialized algorithms and diverse collections of ligand-receptor databases. However, applying existing state-of-the-art tool to newer, high-resolution spatial datasets presents distinct challenges. For instance, our benchmarking revealed that CellNEST, an advanced spatial CCC algorithm developed primarily for lower-resolution platforms like standard Visium, struggled to detect dense, highly localized signaling (e.g., IL-1 encapsulation of abscesses) in high-resolution tissue. This highlights that while such algorithms are powerful, they require significant optimization to fully leverage high-resolution spatial data. TENGU is designed to complement rather than compete with these advanced tools. By prioritizing the generation of mathematically robust segmented units and facilitating their export into open-standard formats (e.g., .h5ad, .zarr), TENGU serves as a high-fidelity gateway. Researchers can utilize TENGU’s CCC module for exploratory analysis, while retaining the flexibility to seamlessly export the cleaned spatial data into more complex, specialized third-party pipelines.

In summary, TENGU offers an adaptable computational framework tailored for the spatial transcriptomic analysis of Visium HD data. By utilizing a transcriptomic-driven cell simulation approach to refine spatial boundaries derived from initial transcript-signal segmentations, the pipeline generates highly cohesive and biologically distinct segmented units. Moving beyond foundational segmentation, TENGU delivers actionable functional insights by seamlessly integrating built-in modules for spatial object embedding, automated high-resolution cell-type annotation, and spatially aware cell-cell communication (CCC) analysis. Together, this integrated approach provides researchers with a robust tool to better characterize and interpret complex spatial dynamics across diverse tissue microenvironments.

## Data availability

The raw data generated by this study is available at NCBI Sequence Read Archive database under BioProject PRJNA1454195

## Software availability

TENGU s publicly available at Github (https://github.com/visanuwan/TENGU)

## Funding

National Institute of General Medical Sciences of the National Institutes of Health (award P20GM125503)

## Bibliography

Visanu Wanchai is an Instructor in the Department of Biomedical Informatics at the University of Arkansas for Medical Sciences. His research focuses on bioinformatics and tool development.

Nancy C. Bustamante-Gomez is a Researcher in the Center for Musculoskeletal Disease Research at the University of Arkansas for Medical Sciences. Her research focuses on the molecular characterization of bone tissue.

Alongkorn Kurilung is a Postdoctoral Fellow in the Department of Biomedical Informatics at the University of Arkansas for Medical Sciences. His research focuses on the experimental development of molecular characterizations and bioinformatics.

Karen E. Beenken is an Associate Professor in the Department of Biochemistry and Molecular Histone Epigenetics at the University of Arkansas for Medical Sciences. Her research focuses on host-microbe interactions.

Sergio Cortes is a Postdoctoral Fellow in the Department of Pharmacology and Toxicology at the University of Arkansas for Medical Sciences. His research focuses on prostate cancer genomics, environmental health, and toxicology.

Mark S. Smeltzer is a Professor in the Departments of Microbiology and Immunology, and Orthopedics at the University of Arkansas for Medical Sciences.

Yuet-Kin Leung is an Associate Professor in the Department of Pharmacology and Toxicology at the University of Arkansas for Medical Sciences. His research focuses on cancer genomics, environmental health, and toxicology.

Jinhu Xiong is an Associate Professor in the Department of Orthopedic Surgery at the University of Arkansas for Medical Sciences. His research focuses on bone biology, specializing in the study of the molecular mechanisms of mechanical loading on bone tissue.

Maria Almeida is a Professor in the Division of Endocrinology and the Department of Orthopedic Surgery at the University of Arkansas for Medical Sciences. Her research focuses on skeletal aging.

Charles A. O’Brien is a Professor in the Division of Endocrinology and the Department of Orthopedic Surgery at the University of Arkansas for Medical Sciences. His research focuses on bone biology, specializing in the study of the molecular regulation of bone tissue by RANKL.

Intawat Nookaew is a Professor in the Department of Biomedical Informatics at the University of Arkansas for Medical Sciences. His research focuses on bioinformatics and applied bioinformatics.

## Notes

### Competing Interest Statement

The authors have declared no competing interest.

### Summary of Updates

Improve the manuscript by modify figure 4 and add figure 5

